# First Efficient Transfection in Choanoflagellates using Cell-Penetrating Peptides

**DOI:** 10.1101/260190

**Authors:** Ruibao Li, Ines Neundorf, Frank Nitsche

## Abstract

Only recently, based on phylogenetic studies choanoflagellates have been confirmed to form the sister group to metazoan. The mechanisms and genes behind the step from single to multicellular organisation and as a consequence the evolution of metazoan multicellularity could not be verified yet, as no reliable and efficient method for transfection of choanoflagellates was available. Here we present cell-penetrating peptides (CPPs) as an alternative to conventional transfection methods. In a series of experiments with the choanoflagellate *Diaphanoeca grandis* we proof for the first time that the use of CPPs is a reliable and highly efficient method for the transfection of choanoflagellates. We were able to silence the silicon transporter gene (SIT) by siRNA, and hence, to suppress the lorica (characteristic siliceous basket) formation. High gene silencing efficiency was determined and measured by light microscope and RT-qPCR. In addition, only low cytotoxic effects of CPP were detected. Our new method allows the reliable and efficient transfection of choanoflagellates, finally enabling us to verify the function of genes, thought to be involved in cell adhesion or cell signaling by silencing them via siRNA. This is a step stone for the research on the origin of multicellularity in metazoans.

## 1. Introduction

Choanoflagellates are a ubiquitous distributed group of single-celled microeukaryotes, which are able to form colonies. They are characterized by a unique morphology inlcuding a protoplast with a single flagellum surrounded by a collar of microtubule. Choanoflagellates play an essential role in the microbial food web as they are highly efficient filter feeder, mainly feeding on bacteria (Leadbeater, 2015). Interest in the evolutionary biology of choanoflagellates has increased, since their position as the closest sister-group to metazoan in the eukaryotic group Opisthokonta has been verified (Adl et al., 2012; Carr et al., 2008; Ruiz-Trillo et al., 2008). The last common ancestor of metazoan and choanoflagellates likely originated from the same ancestral choanoflagellate (Carr et al., 2008; Nielsen, 2008; Steenkamp et al., 2006). Choanoflagellates possess several genes also found in metazoans, which are involved in cellcell communication and transfer (Fairclough et al., 2013; King and Carroll, 2001; King et al., 2008).The function of these genes in choanoflagellates remains untested as no gene-silencing method for these protists is available. Especially with respect to the evolution of multicellularity, gene silencing is an essential tool to understand and proof the role of candidate genes. All conventional methods like micro-injection, electroporation, freeze-thaw techniques, viral delivery systems, liposomes or cationic lipids, are not efficient and reliable methods for transfection of choanoflagellates until now. Here, we present cell-penetrating peptides (CPPs) as an alternative.

CPPs have been applied as a useful carrier to deliver cargoes into metazoan cells over the last two decades (Crombez et al., 2009; Dai et al., 2011; Frankel and Pabo, 1988; Green and Loewenstein, 1988; Hoyer and Neundorf, 2012; Keller et al., 2014; Lundberg et al., 2007; Rennert et al., 2008; Neundorf et al. 2009; Säälik et al., 2004). For the first time we proof that transfection of the choanoflagellate *Diaphanoeca grandis* by CPP mediated siRNA is a reliable and high efficient method. This method opens new opportunities to investigate the evolution of multicellularity by knocking down candidate genes.

## 2. Materials and Methods

### 2.1 Materials

N^α^-Fmoc protected amino acids were purchased from IRIS Biotech (Marktredwitz, Germany). Side chains of the trifunctional amino acids were protected using acid-labile protecting groups: Pbf for Arg; Trt for Gln and His; Boc for Lys; *tert-Butyl* for Glu and Thr. For introducing the side chain Fmoc-Lys(Dde)-OH was used.

For gene silencing, silicon transporter gene alpha (SITalpha) siRNA (Eurofins, Germany: sense 5’AACAAUGGAACAACCCUCCAU) and scrambled siRNA (Eurofins, Germany: sense 5’CCUUGAUUGGUUGUUGGGA) for negative control were designed based on a study of the SIT in choanoflagellates, NCBI accession no. KU821737 (Marron et al., 2013).

### 2.2 Peptide synthesis

Peptides used were synthesized by automated peptide synthesis on a multiple peptide synthesizer (MultiSyntech, Syro) via fluorenylmethoxycarbonyl (Fmoc)/ter-butyl (t-Bu) strategy according to ref. Neundorf et al 2009. In brief, the sequence FHTFPQTAIGVGAP was synthesized on Rink amide resin (loading was 0.5 mmol/per gram resin) using Fmoc-protected amino acids (7 eq) and Oxyma Pure^®^/*N’,N*-diisopropylcarbodiimide (DIC) (7 eq) activation in DMF. Fmoc-Lys(Dde)-OH was introduced manually by the same activation procedure. The peptide was then elongated using the peptide synthesizer to Boc(butyloxycarbonyl)- GLLEALAELLEE—K(Dde)-FHTFPQTAIGVGAP. The Dde group was cleaved by treating the resin with a hydrazine/DMF solution (3%) at least 10 times for 5 min. Then the side sequence was synthesized (KKRKAPKKKRKFA) using automated peptide synthesis and Oxyma Pure^®^/DIC activation as described above. For uptake studies, the peptide was additionally labeled with 5, 6-carboxyfluorescein (using 5 eq HATU (O-(7-azabenzotriazole-1-yl)-N,N,N’,N- tetramethyluronium-hexafluorophosphate) and 5 eq DIPEA (diisopropylethylamine)) at the N- terminal of the side sequence.

After complete synthesis the peptides were cleaved from the resin using trifluoroacetic acid (TFA) and triisopropylsilane as scavenger (90:10 v/v), and precipitated in ice cold diethyl ether. Purification was achieved by preparative RP-HPLC on a Hitachi Elite LaChrom instrument (VWR, Darmstadt, Germany) at 6 ml/min flow rate and 220 nm detection. The chromatography was performed on a 15 x 250 mm Jupiter 4u Proteo 90A column (Phenomenex, Aschaffenburg, Germany) by using a linear gradient of 10−60% B in A (A = 0.1% TFA in water; B = 0.08% TFA in ACN) in 45 min at 6 mL/min. Detection was at 220 nm wavelength and the fractions containing the peptide were collected. Solvents were removed under reduced pressure and the peptide was lyophilized overnight. Purity was >95%. Peptides were analyzed by LC-MS using an Agilent HPLC instrument with parallel detection at 220 nm UV-absorption and electrospray-ionization mass spectrometry (ESI-MS, Finnigan MAT, Thermo). The reversed-phase analytical HPLC (RP-HPLC) was performed at 1.2 mL/min flow rate on a 4.6 x 100 mm Kinetex 2.6u C18 100A column (Phenomenex, Aschaffenburg, Germany). A gradients of 10 - 60 % acetonitrile in water over 15 min (with constant 0.1 % TFA) was used: N-E5L-hCT(18−32)-k7 (CPP): MW_calc_. 4317.6 Da, MW_exp_. 4317.9 Da, [M+5H]^5+^: 864.71 m/z, [M+4H]^4+^: 1080.49 m/z, [M+3H]^3+^: 1440.13 m/z; N-E5L-hCT(18-32)-k7(CF) (CF-CPP): MW_calc_. 4675.3 Da, MW_exp_. 4677.6 Da, [M+5H]^5+^: 936.82 m/z, [M+6H]^6+^: 780.72 m/z, [M+7H]^7+^: 669.22 m/z.

### 2.3 Cultivation and growth rate of Diaphanoeca grandis

*D. grandis* was cultivated in artificial seawater (ASW), 25 practical salinity units (PSU) at 13°C in 50ml culture flasks (Sarstedt, Nümbrecht Germany). To maintain constant conditions and to exclude bacteria as an interfering factor for the experiments, heat-killed bacteria *(Pseudomonas pudita)* were added as food source at a volume ratio of 20:1 every 72 hours in addition to the natural bacteria community. Growth rate was determined by mini automated cell counter (Moxi ORFLO, VWR, Germany) for 16 days with daily measurements of three replicates.

### 2.4 Cell viability test and Internalization of CPP by Diaphanoeca grandis

Different concentrations of CPP (10μM, 20μM, 30μM, 40μM, 50μM, 60μM and 70μM) were tested on *D. grandis*, to find the optimal, nontoxic concentration. After incubating for three and six hours, cells were stained by DAPI (3μg/ml) and PI (0.6μg/ml) for 10 min for live/dead staining (Andras et al., 2000). Stainded cells were washed with ASW five times, and then counted by a Keyence inverted fluorescence microscope using a 40× objective with phase contrast and appropriate filters For positive controlls, cells were treated with 70% alcohol for 10 min at 13°C, for negative controls cells were only washed with ASW. For internalization experiments, triplicates of *D. grandis* from the same clonal culture were inoculated in 8-well tissue culture chambers (GmbH, Sarstedt Germany) and treated with 30μM CF-CPP at 13°C for 1h to observe the internalization process Cells were washed five times with ASW medium carefully, and then images were captured with a Keyence inverted fluorescence microscope using a 40× objective with phase contrast and additional green fluerscent filter set (λex=489, λem=509 nm). As a control, three parallels of untreated *D. grandis* were observed under the same conditions.

### 2.5 Electromobility shift assay (EMSA)

In order to find the optimal chare ratio between CPP and siRNA, an electromobility shift assay was performed as in Hoyer and Neundorf, 2012. In brief, siRNA (100pmol/μl) was incubated at 37°C for 60 min with CPP stock solution (1mM) to form the complexes at charge ratios of 2:1, 5:1, 10:1, 15:1, 20:1, 30:1 and 40:1 (CPP/siRNA) using ASW for dilution to generate the same conditions as in the transfection experiments. Afterwards, these mixtures were analyzed by electrophoresis for 1.5h at 140 V on a 1.5% agarose gel stained by ethidium bromide in tris-acetate-EDTA (TEA) buffer. Pure siRNA was used as control.

### 2.6 CPP-mediated transfection of siRNA

1μl of 100μM siRNA were mixed with 4.8 μl of 1mM CPP stock solution(charge ratio of 10:1 (CPP/siRNA)) and incubated at 37°C for 60 min. CPP/siRNA complexes were added to cells with ASW (the final concentration of CPP was 30 μM) at 13°C for 6 hours for transfection. After two days cultures were inspected using inverted light microscopy, and cells having a complete, partial and no lorica were counted. All experiments were done at least in triplicates and repeated three times.

### 2.7 Electron microscopy

50 hours after transfection cells were fixed with glutardialdehyde in cacodylate buffer (final concentration 3% and 0.05M, respectively) at 4°C for 30 min. Post fixation was done using osmium tetroxide (OsO_4_) for 10 min, followed by dehydration in an ethanol series (30%, 50%, 70%, 80%, 90% and 97% ethanol for 10 min). For final dehydration cells were treated with a 1:1 solution of hexamethyldisilazane (HMDS) and 97% ethanol for 15min and then transferred to 100% HMDS for 10min. After removing HMDS, cells were allowed to fall dry at room temperature. The samples were sputter coated with a 120Å layer of gold and observed using a FEI Quanta 250 FEG.

### 2.8 RNA isolation and real-time quantitative PCR

Total RNA was extracted with pepGOLD total RNA kit, and cDNA was synthesized using a Biozym cDNA synthesis kit with oligo primer, according to the manufacturer’s recommendation. 75ng of cDNA was used for real-time quantitative PCR (RT-qPCR) followed the Biozym Blue S’Green qPCR kit. The primer was designed as 5’-AAGTTCACCTACTGGTGGTCGT-3’ (forward) and 5’-GCATCACAATGAAGAGGCAGATAT-3’ (reverse) from Eurofins (Germany). Reactions were analyzed upon an Applied Biosystems StepOne ^TM^ Real-Time PCR system using the following cycle conditions: 95°C for 2 min., followed by 40 cycles at 95°C for 5 sec. and 60°C for 25 sec. The conditions of met stage are 95°C for 15 sec, 61°C for 1min. and 95°C for 15 sec. Data was analyzed with StepOne software V2.0.

### 2.9 Statistical calculation

Results were showed as means ± standard deviations. Mean values and standard deviations were calculated for each sample (triplicates) examined in at least three independent experiments. The level of statistical significance was set at P<0.05 and detected by ANOVA.

## 3. Results

### 3.1 Cytotoxicity and Internalization

*Diaphanoeca grandis* (20ml) was cultivated in 50ml culture flasks at 13°C for 16 days and showed an exponential growth phase, which is optimal for transfection after seven days (Fig. 1A).The optimal time point for transfection was determined to be between seven and nine days of growth under these conditions.

**Figure 1.**
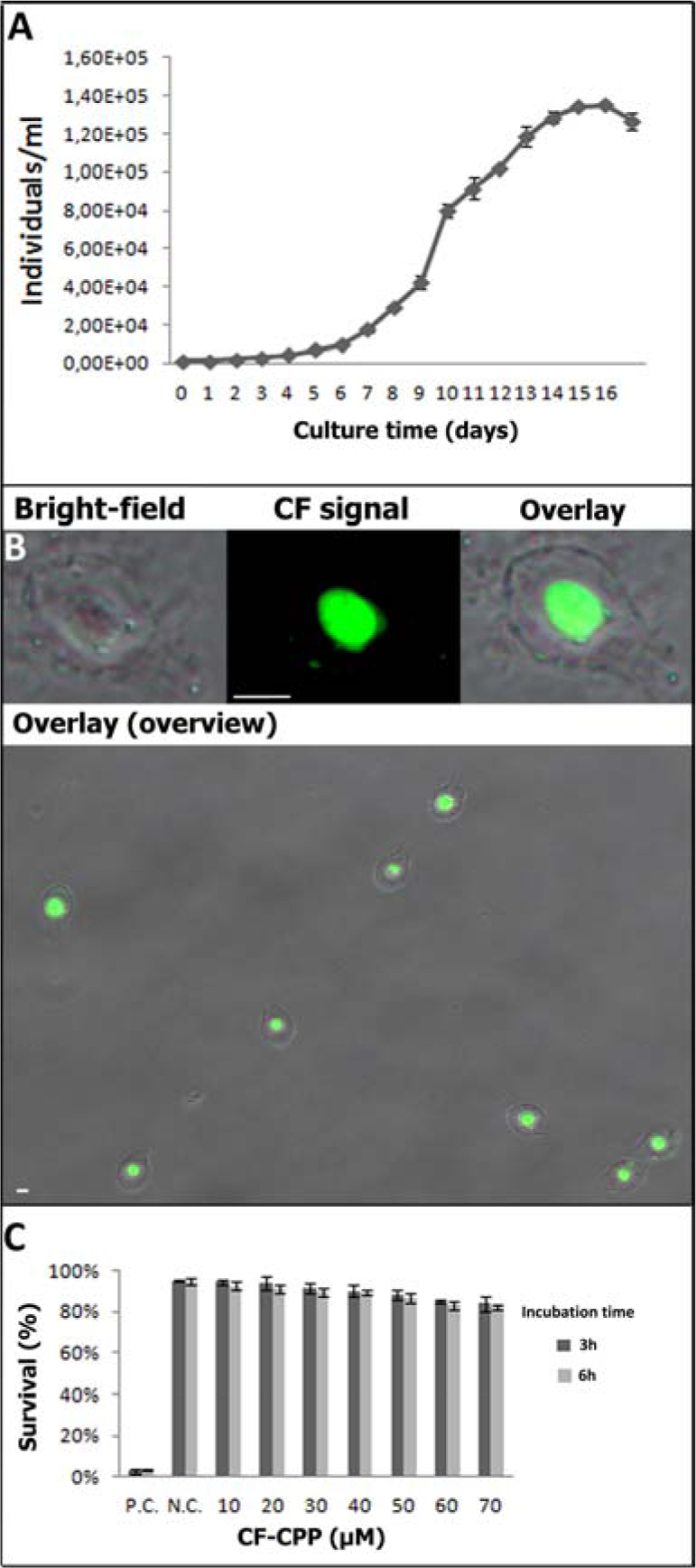
Growthrate and internalization and toxicity. A: Growth rate of *D. grandis.* B: Uptake of CF-labeled CPP in *D. grandis.* Cells were inspected using light and fluorescence microscopy. The whole protoplast is evenly stained by the CPP. Scale bar is 5|jm. C: CPP influence on cell viability. Different concentrations of CF- CPP were incubated with *D. grandis* for 3h or 6h, respectively. P.C.= positive control (70% alcohol) and N.C.= negative control, respectively.

The CPP N-E5L-hCT(18-32)-k7 used in this study (for more details see Materials and Methods) was recently developed by us for efficient siRNA delivery into mammalian cells (Hoyer and Neundorf, 2012). It consists of a branched structure that was designed to support complexation of the positively charged peptides with the negatively charged oligonucleotides (Rennert et al. 2008). Furthermore, the sequence N-E5L was attached aiming to increase the cytosolic uptake (Neundorf et al. 2009). First we tested its influence on cytotoxicity in order to find the optimal concentration for transfection in *D. grandis.* Therefore, the cells were treated with a broad range of CPP concentrations and two different periods. Afterwards, cells were washed and live cells were counted. We observed that the CPP showed almost no toxicity to *D. grandis* even when incubating the cells with 70μM solutions for 3h and 6h (84% ± 2%, 82% ± 1% survival), respectively (Fig. 1C). to In a next set of experiments we elucidated the peptide uptake in *D. grandis.* For this, different concentrations of the CF-labeled CPP were incubated with the cells at 13°C. We observed that a peptide concentration of 30μM resulted in high efficient cellular uptake. Within one hour after treatment strong fluorescence in *D. grandis* was visible that was distributed throughout the whole protoplasm (Fig.1 B). Notably, we exclude an uptake of the CPP via bacteria. Since most food bacteria where heat-killed bacteria, the number of internalized viable bacteria was too low to result in the intensity shown in Fig.1B.

### 3.2 Transfection studies

For further gene silencing studies, we aimed to transport siRNA into *D. grandis.* We chose to silence the gene for the silicon transporter (SIT), which is responsible for the formation of the lorica (Marron et al., 2013). First, we investigated the complex formation between peptides and siRNA byperforming an electro mobility shift assay (EMSA). Different peptide/siRNA charge ratios were tested ranging from 1:1, 5:1, 10:1, 15:1. Already at a charge ratio of 5:1 peptide/siRNA, the nucleic acids stayed inside the gel pockets pointing to strong binding of the peptides to the siRNA (Figure S1). However, a charge ratio of 10:1 was used for further studies to assure a proper complexation.

Next we proved CPP-mediated gene silencing in *D. grandis* by transfecting the organisms with CPP/siRNA complexes. As negative controls untreated cultures, CPP only, SIT siRNA only, and CPP + scrambled siRNA (Fig. 2A) were used to exclude biased signals. After two days, the cells were analyzed with respect to a proper lorica formation. Interestingly, only the transfection treatments with CPP/siRNA complexes revealed naked cells without the typical silica basket (Fig. 2A, Fig. 3A). The knockdown efficiency was 68% ± 2% (Fig. 2B), as calculated from the percentage of transfected naked cells or cells with inclompete lorica (S2) over the total cell numbers. In addition, a series of time course (2h, 6h, 12h and 50h) experiments were set up to quantify changes in SIT mRNA expression level. The transcript abundance of SIT was suppressed to 27%± 3% from 2h and to8%±2% after 12h but recovered after 50h post-treatment to 15%±2% (Fig. 3B). Thus, we concluded that SIT mRNA expression was highly significantly suppressed in *D. grandis* after CPP-mediated SIT-siRNA delivery.

**Figure 2.**
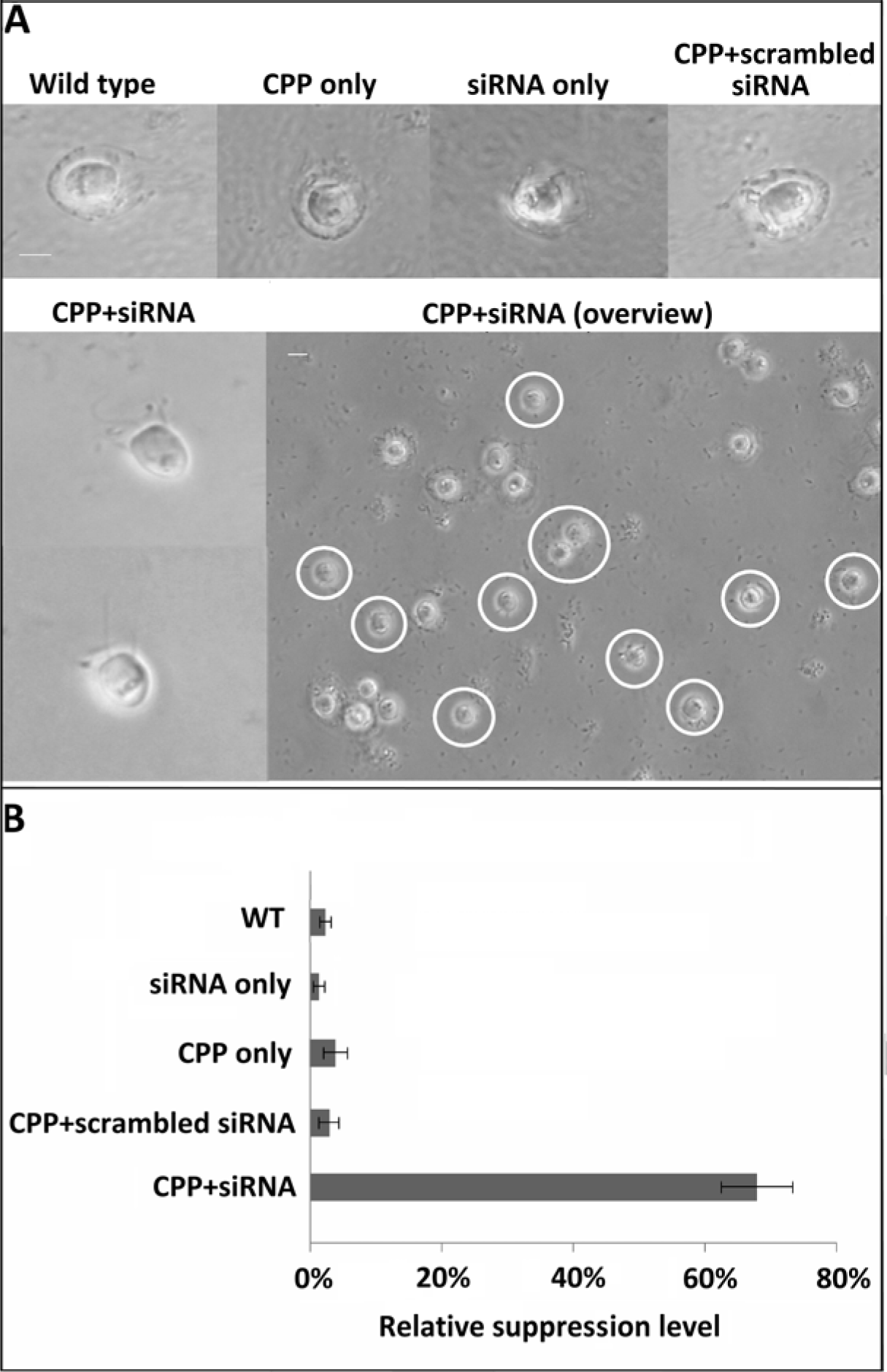
Transfection efficiency. A: 50h after transfection light microscopy images of *D. grandis* were taken. As control, cells treated with ASW medium only, CPP+ scrambled siRNA, siRNA only or CPP only are shown. Cells treated with 30μM/ 1.2μM of CPP/SIT- siRNA complex were naked after 50h (white circles). Scale bar is 5μm. B: Relative suppression of SIT gene expression was determined by counting naked cells.

Transfected cells were unable to produce the costal strips, which are normally accumulated prior to cell division. As a result, up to 50h post transfection a next generation evolved that lacked a lorica (Fig. 2A, Fig. 3A). After the siRNA was degraded and no further gene silencing possible, the naked, lorica-less generation started to accumulate costal strips for the third generation, which then again produced a full lorica (Fig. 3A). The naked generation remained without lorica, as tectiform acanthoecids are not able to produce their own lorica (Leadbeater 2015).

**Figure 3.**
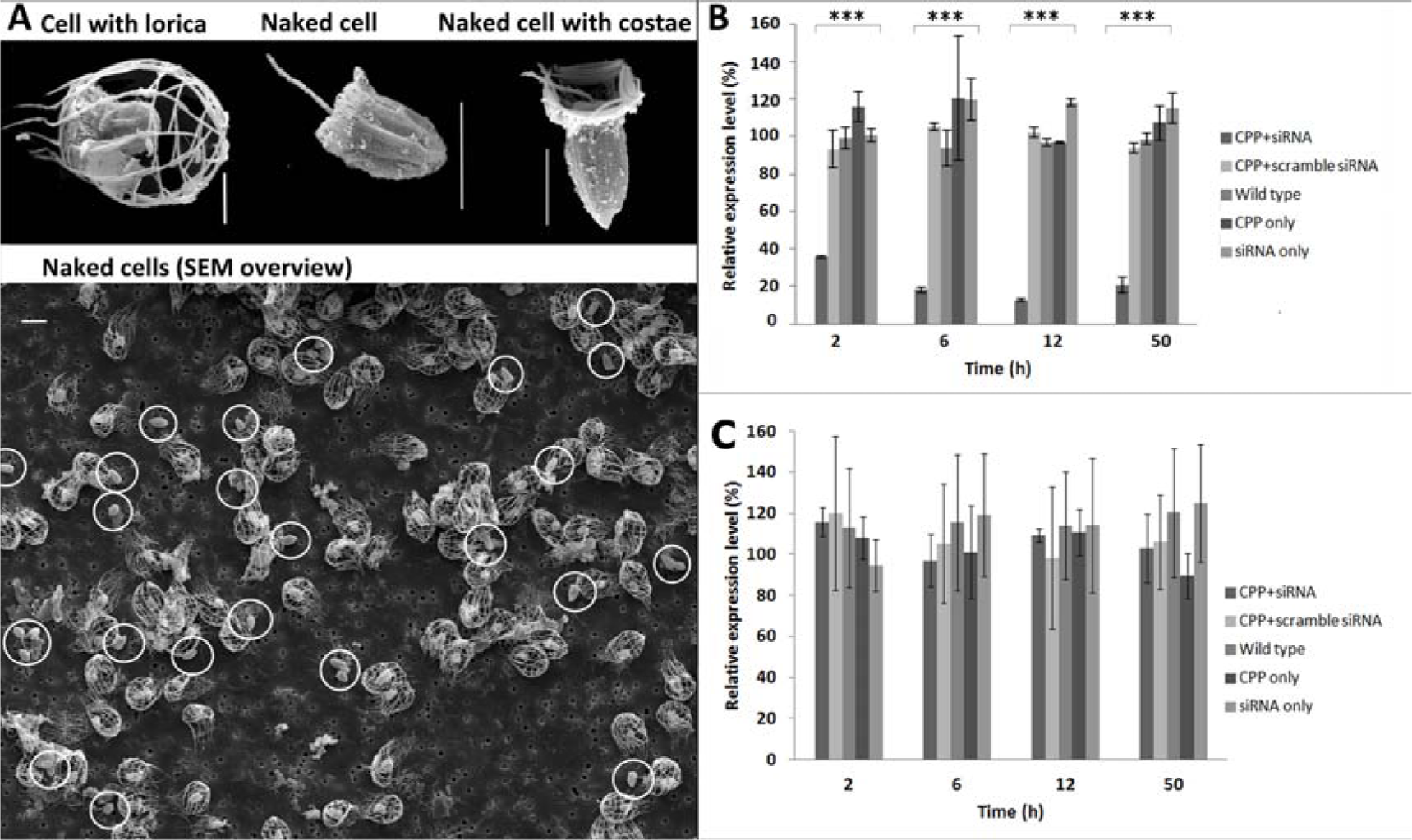
Reproduction after transfection and qPCR A: 50 h after transfection, electron microscopy images of *D. grandis* were taken. The second, transfected generation of loricated mother cells was naked; those naked cells produced costal strips for their next generation, which were loricated afterwards. Naked cells are marked with circles. Scale bar is 5μm. B: Real-time quantitative PCR analysis of SIT gene expression in *D. grandis.* Samples were taken at different time points (2h, 6h, 12h, and 50h). ANOVA, n=3, *P < 0.05, **P < 0.01, ***P < 0.001.

## 4. Discussion

Since the last two decades cell-penetrating peptides emerged as a successful method to deliver cargos into target cells. However, to the best of our knowledge, this is the first time that CPPs were used to transport siRNA into a choanoflagellate species. Other transfection methods that utilizeliposomes, micro-injection or electroporation all failed, since they were either toxic or lethal to choanoflagellates, or showed only low transfection efficiency. This lack of appropriate transfection tools is the reason that research on the evolution of multicellularity is still hindered, and that, for example, the function of candidate genes previously found regarding cell adhesion, cell signaling, and colony formation, is still not verified. Although the CPP tested in this study (namely N-E5L-hCT-(18-32)-k7) was recently developed for mammalian siRNA transfection, it seemed to be an appropriate tool for transfection of choanoflagellates, too. To prove the function and efficiency of this method, we silenced a non-housekeeping gene in *D. grandis*, responsible for a clear morphologic character, the lorica. The silicon transporter (SIT) gene was considered as the appropriate candidate gene for our experiment, sinceit has a clear function and silencing is not lethal̤Silicon is the second abundant element after oxygen in the Earth’s crust, soil and nearly all waters. In *D. grandis* silicon is essential to build up the lorica, a species specific morphologic characteristic. This basket like structure is composed from single siliceous rods, the costae. During reproduction the mother cell incorporates silicon from the exterior by concentrating silicon with the silicon deposition vesicle to produce a complete set of costae for the next generation (Marron et al., 2013). In our experiments we show that silencing the SIT gene in *D. grandis* results in a next generation with either no, or incomplete lorica, depending on the production stage the mother cell was in during transfection. This generation will, after the siRNA is degraded, return to produce costae for the third generation, whereas the second generation will stay naked (Fig. 2A, Fig. 3A). One of the major limitations of non-viral gene delivery systems to deliver cargoes into cells is poor release from the endosomal compartment after internalization. It is a long journey for siRNA to silence target genes from crossing cell membrane to degrade mRNA. However, after 2h already 72% of SIT mRNA was suppressed after interfering by siRNA transferred by the CPP (Fig.3B). This indicates a rapid release of siRNA into the cytoplasm to form the RISC and to promote down-regulation of SIT mRNA robustly and efficiently.

It has been already discussed that cytotoxic effects of CPPs might affect cell membrane integrity and viability negatively (Niles et al., 2007; Wu et al., 2007). The amount of CPPs and/or CPP- cargo aggregates used is limited depending on cytotoxic effects that occur during the internalization and transfection process. Many factors, such as incubation time, cargoes, used CPPs, the concentration of CPPs, and ions might affect cytotoxicity (Andaloussi et al., 2007; Jones et al., 2005; Saar et al., 2005; Sasaki et al., 2008). Therefore, it is highly necessary to establish optimal conditions for every distinct transfection experiment using CPPs. These conditions might even vary from species to species and strongly depend on the salinity of the media used for culturing.

Preliminaryexperiments using plasmid DNA (eGFP) with eukaryotic specific promoters for transfection did not result in an expression of the plasmid DNA in choanoflagellates only, but also in bacteria, although they lack the specific promotor (data not shown). In future, we plan to insert? choanoflagellate specific promoters in a fluorescent plasmid, with the aim to label choanoflagellates permanently.

Within this study we present for the first time a suitable and high efficient method for transfection of choanoflagellates. The fact that within the two model organisms of choanoflagellates, *Monosiga brevicollis* and *Salpingoeca rosetta*, no complete RNAi pathway was detected, hinders the usage of siRNA to silence candidate genes. We plan to use the newly developed gene editing method CRISPR/Cas9 in combination with our CPP to overcome this obstacle. However, for several other choanoflagellate species, main components (Argonaute and Dicer) of this pathway were found (Richter, 2013). This opens a new door for studies on the evolution of multicellularity.

## Acknowledgements

We thank A. Jeuck for establishing RT-qPCR and A. Klimpel for providing the CPP. Special thanks to Prof. Wostemeyer (University of Jena), D. Richter (Station Biologique de Roscoff) and N. King (University of Berkely) for fruitful discussions. Many thanks to H. Arndt for offering labspace and support.

---

**Figure S1.**
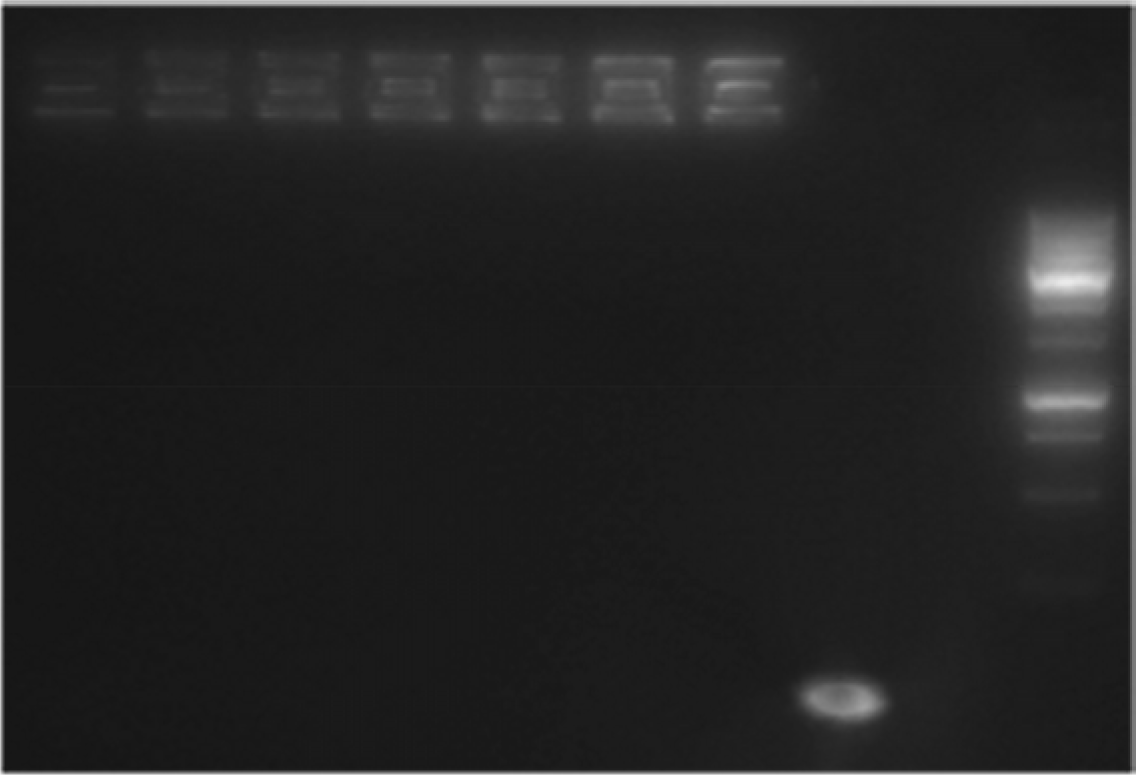
Electromobility shift assay of the interaction between CPP and siRNA (different charge ratios peptide/siRNA are shown). M = DNA ladder (Genaxxon bioscience), C= negative control, CPP only.

